# The *Arabidopsis* amino acid transporter UmamiT20 confers *Botrytis cinerea* susceptibility

**DOI:** 10.1101/2024.10.26.620370

**Authors:** Matthew J. Prior, Diana Weidauer, Jui-Yu Liao, Keiko Kuwata, Federica Locci, Chen Deng, Hong Bo Ye, Qiang Cai, Margot Bezrutczyk, Chengsong Zhao, Li-Qing Chen, Martin C. Jonikas, Guillaume Pilot, Hailing Jin, Jane Parker, Wolf B. Frommer, Ji-Yun Kim

## Abstract

Induction of SWEET sugar transporters by bacterial pathogens via transcription activator-like (TAL) effectors is necessary for successful blight infection of rice, cassava and cotton, - likely providing sugars for bacterial propagation.
Here, we show that infection of *Arabidopsis* by the necrotrophic fungus *Botrytis cinerea* causes increased accumulation of amino acid transporter UmamiT20 mRNA in leaves. UmamiT20 protein accumulates in leaf veins surrounding the lesions after infection. Consistent with a role during infection, *umamiT20 knock-out* mutants were less susceptible to *B. cinerea*.
Functional assays demonstrate that UmamiT20 mediates amino acid transport of a wide range of amino acid substrates.
Pathogen-induced UmamiT20 mRNA and protein accumulation support the hypothesis that transporter-mediated pathogen susceptibility is not unique to SWEETs in bacterial blight of rice but also for a necrotrophic fungus and implicate nutrients other than sucrose, i.e., amino acids, in nutrition or nutrient signaling related to immunity. We hypothesize that stacking of mutations in different types of susceptibility-related nutrient carriers to interfere with access to several nutrients may enable engineering robust pathogen resistance in a wide range of plant-pathogen systems.

**Lay Abstract:** Pathogens infect plants to gain access to their nutrient resources, enabling the pathogens to cause disease and reproduce efficiently. Here we find that an amino acid transporter constitutes a susceptibility factor for the fungal pathogen *B. cinerea*.

## Introduction

Plant pathogens cause substantial yield losses in agriculture; therefore, the engineering of resistant crops is of utmost relevance. Several concepts for introducing resistance have been developed, e.g. transfer of pattern recognition receptors from different plant species and mutation of susceptibility genes (Schwessinger *et al*., 2015; Blanvillain-Baufumé *et al*., 2017). *Botrytis cinerea* (*B. cinerea*) is a necrotrophic pathogen that infects vegetables, flowers, and fruits including grapevine. Depending on the conditions, *B. cinerea* causes grey mold or bunch rot, which can seriously damage grape yield and quality. *B. cinerea* can be utilized to cause noble rot, thereby increasing positive traits for vinification (Blanco-Ulate *et al*., 2015). Fungicides can be used to control infections. However they are costly, may have a negative perception by consumers, may be harmful to animals and humans, and resistance against various fungicides continues to emerge (Kim *et al*., 2016).

Pathogens require host nutrients for efficient propagation, and it has been suggested that solute efflux from host cells may exert control over the transfer of solutes from host to pathogen (Patrick 1989). Recent work indicates that pathogens induce host transporters to gain access to host nutrient resources as a virulence mechanism. SWEET sugar transporters play critical roles in sugar transport, including phloem loading, nectar secretion, seed filling, microbiota colonization, as well as pathogen susceptibility (Chen *et al*., 2010, 2015; Li *et al*., 2012; Lin *et al*., 2014; Cohn *et al*., 2014; Zhou *et al*., 2015; Loo *et al*., 2024). Engineering of the pathways that lead to pathogen-triggered activation of transporters may enable limiting access to host nutrients. An example is the successful implementation of resistance via genome editing of the binding sites of bacterial effectors in host *SWEET* uniporter gene promoters (Eom *et al*., 2019; Oliva *et al*., 2019; Schepler-Luu *et al*., 2023). *Xanthomonas* uses transcription activator-like (TAL) effector proteins bind to host promoters, thereby triggering activation of *SWEET* gene transcription (Chen *et al*., 2010). Surgical CRISPR or TALEN-based mutagenesis of the promoter sequences to which the effectors bind abolishes TAL activation of host genes, thus preventing SWEET transporter induction and causing resistance (Li *et al*., 2012; Bezrutczyk *et al*., 2018a; Eom *et al*., 2019; Oliva *et al*., 2019; Wu *et al*., 2022; Schepler-Luu *et al*., 2023). SWEETs have been shown play important roles in other pathosystems as well (Chen *et al*., 2023).

VvSWEET4 had been identified as a susceptibility factor of grapevine to *B. cinerea* (Chong *et al*., 2014). *B. cinerea* infection of the *Arabidopsis sweet4* knock-out mutants resulted in reduced disease symptoms (Chong *et al*., 2014). If we hypothesize that pathogens infect plants primarily to gain access to host nutrients for reproduction, one may propose that any essential nutrient could be limiting (Liebig’s hypothesis), or be made limiting by the host (van der Ploeg *et al*., 1999; Bezrutczyk *et al*., 2018b). Based on this hypothesis, we surmise the pathogens dependence on access to a full suite of essential micro- and macroelements from the host. Hence transporters for organic nitrogen may also be candidates for pathogen susceptibility. UmamiTs, transporters which can function in amino acid efflux to play roles in the translocation of organic nitrogen from leaves to seeds, analogous to the role of SWEETs in sugar allocation (Müller *et al*., 2015; Zhao *et al*., 2021). Analysis of public data bases indicated that the mRNA levels of *UmamiT20* increased during *B. cinerea* infection of *Arabidopsis*. We here show that translational UmamiT20-GFP fusions localized predominantly to the plasma membrane. Functional assays showed that UmamiT20 can mediate the cellular export of a broad range of amino acids when expressed in *Xenopus* oocytes. During *B. cinerea* infection, UmamiT20-GUS translational fusion proteins accumulated in the vasculature surrounding the infection sites, and *umamit20 knock-out* mutants showed decreased susceptibility to *B. cinerea* infection. Together this work implicates a member of the UmamiT transporter family as an *Arabidopsis* susceptibility gene for *B. cinerea* infection, possibly expanding the concept of nutrient transport as a susceptibility factor from carbon to nitrogen, from biotrophs to necrotrophs, and from bacteria to fungi.

## Materials and Methods

### Bioinformatic analyses

Online microarray or RNA-seq data was accessed from referenced publications and analyzed in Microsoft Excel to determine gene expression changes during infection. Publications and GEO datasets accessed are available in Table 1.

**Table 1.** Compilation of microarray or RNA-seq data for accumulation of mRNAs for selected transporter genes (*SWEET, UmamiT*, and *STP)* during *B. cinerea* infection of *Arabidopsis* leaves using different infection protocols and the different inoculation buffers for fungal spores during the infection assays. The time point in hours (h) with the maximal -fold transcript accumulation (≥ 2-fold) or decrease (≤ 0.5-fold) is presented. No significant change is indicated by “nsc”, while “n/a” indicates the transcript levels were not measured in that study. For the GSE48207 study the Col-0 T0 time point was used as a control as there was no mock infected time point available for reference.

| Study | GSE ID | Inoculation buffer | *B. cinerea* isolate | Time points | *SWEET4* | *UmamiT18* | *UmamiT20* | *STP13* |
| --- | --- | --- | --- | --- | --- | --- | --- | --- |
| Birkenbihl, 2012 | no GSE ID | potato dextrose | CECT2100 | 14 h | n/a | 1.58 at 14 h | nsc | 29x at 14 h |
| Coolen, 2016 | RNA-seq | potato dextrose | B05.10 | 3, 6, 12, 18, 24 h | nsc | nsc | nsc | 5.6x at 18 h |
| Ferrari, 2007 | GSE 5684 | potato dextrose | SG1 | 18, 48 h | nsc | 3x at 48 h | 4x 48 h | 15x at 48 h |
| Ingle, 2015 | GSE 70137 | grape juice | Pepper | 18, 22 h | nsc | 2x at 22 h | nsc | 2x at 22 h |
| Lemmonnier, 2014 | no GSE ID | potato dextrose | B05.10 | 48 h | n/a | n/a | n/a | 6x at 48 h |
| Mulema, 2012 | GSE 24445 | grape juice | Pepper | 12, 24 h | nsc | nsc | nsc | 3x at 12 h |
| Windram, 2012 | GSE 39598 | grape juice | Pepper | 2-48 h | 5x @ 38 h | 6x at 34 h | 2x 36 h | 8x at 30 h |
| Zhang, 2013 | GSE 48207 | potato dextrose | B05.10 | 0-48 h | nsc | 0.5x at 6h | nsc | nsc |

### Plant growth conditions

Seeds were sown onto ½ salt strength Murashige Skoog (MS) media, supplemented with 1% sucrose and grown for 1 week at a Photosynthetic Photon Flux Density (PPDF) of 120 µmol m^-2^ s^-1^ in a growth cabinet. After one week, seedlings were transferred to soil. Plants were grown in 10-hour light/14-hour dark conditions at 22 °C when the lights were on (∼120 µmol m^-2^ s^-1^), and 21 °C when lights were off, and watered twice every week. Fertilizer was not used for plant growth. Fully extended rosette leaves were infected by drop inoculation after 5 weeks of growth. Plants with leaves that showed signs of damage were excluded from the experiments.

### Transient gene expression in *Nicotiana benthamiana* leaves

The UmamiT20 ORF was amplified by PCR using the primers pair *UmamiT20* ORF fw and *UmamiT20* ORF rev (Table S1) using reverse transcribed *Arabidopsis* RNAs. The resulting PCR amplicon was then cloned into pDONR™/Zeo (Gateway™ vector from Invitrogen, MA, USA), validated by sequencing, then mobilized into the binary expression vector pAB117. The *Agrobacterium tumefaciens* (*A. tumefaciens*) strain GV3101-p19 MP90 was transformed using the binary expression vector pAB117 carrying the *UmamiT20* ORF and a C-terminally fused enhanced GFP (eGFP) under control of the ß-estradiol inducible XVE transactivator-based promoter (Somssich *et al*., 2015) or transformed with binary expression vector pAB118 carrying *ZmSWEET13a* C-terminally fused with mCherry driven by a *Cauliflower mosaic virus* 35S promoter (Bezrutczyk *et al*., 2018a). *Agrobacterium* culture and tobacco leaf infiltration were performed as described (Sosso *et al*., 2015). Chloroplast fluorescence was detected on a Zeiss LSM 780 or 880 confocal microscope (470 nm excitation with simultaneous detection from 522–572 nm (eGFP), 561 nm excitation with detection from 600-625 nm (mCherry) and 667-773 nm detection of chloroplast fluorescence. Image analysis was performed using Fiji (https://fiji.sc/) and Omero software (Oliva *et al*., 2019).

### Generation of *umamiT20-2* mutant using the CRISPR-Cas9 system

Target sequences were designed at the first exon of *UmamiT20* using CHOPCHOP (http://chopchop.cbu.uib.no) and CRISPOR (http://crispor.tefor.net). Primers DW185 and DW186 (Table S1) which include the target sequences flanked by the *Bsa*I site were used for amplifying the gRNA scaffold and U3b promoter originating from pENTR4-sgRNAs backbone-based vector (Zheng *et al*., 2020). Amplified amplicons were purified (Macherey-Nagel, Düren, Germany) and assembled in the pAGM55261 (Addgene) vector using the Goldengate method, as described in the NEB Golden Gate Assembly protocol (5-10 inserts protocol) (NEB, MA, United States). *E. Coli* Top10 competent cells (One Shot TOP10 chemically competent, Thermo Fisher Scientific, Darmstadt, Germany) were then transformed using the assembled product. Positive clones were selected by colony PCR using primer pairs MR9 + DW186 (Table S1) and confirmed through restriction enzyme digest (*EcoRV-HF*/*PmeI*) and sequencing. The electrocompetent *A. tumefaciens* (GV3101) were transformed using the binary construct. Col-0 plants were used for floral dipping (Clough & Bent, 1998). For mutant characterization, T_1_ transgenic plants were selected on ½ strength MS medium supplemented with glufosinate ammonium (10 µg/ml) and cefotaxime (100 µg/ml). Mutations in *UmamiT20* were examined by amplifying the region-of-interest using gene-specific primers MQ5fw and MQ5rev followed by Sanger sequencing (primer MQ5 seq; Table S1). Cas9-free plants were selected by genotyping and screening for the lines absent of the seed-coat specific RFP fluorescence. Primers sequences are available in Table S1.

### Transport assays in *Xenopus* oocyte

*UmamiT20* ORF cloned into pDONR™/Zeo (Gateway™ vector from Invitrogen, MA, USA) was mobilized into the oocyte expression vector p002-GW by LR reaction (Invitrogen, MA, USA). The pOO2 plasmid was linearized using *Mlu*I restriction enzyme and UmamiT20 cRNA was synthesized using the mMessage Machine SP6 Kit (Invitrogen, MA, USA) as described in Chen *et al*., 2010. Oocytes were purchased from Ecocyte Biosciences. 50 nl of *UmamiT20* cRNAs (>2ng/µL) or RNAse-free water, a standard control for oocyte experiments, was injected into oocytes. Oocytes were incubated in ND96 solution supplemented with 100 µM gentamycin at 16 °C for 2 days to allow for protein synthesis. For efflux assays, oocytes were injected with 50 nl ^15^N amino acid mixture (catalog number 767972, Sigma Aldrich, MO, USA) and immediately placed in ice-cold ND96 solution for 10 min to allow closure of the injection spot. The ^15^N amino acid mixture consisted of aspartic acid (60 mM), threonine (35 mM), serine (35 mM), glutamic acid (40 mM), proline (20 mM), glycine (100 mM), alanine (100 mM), valine (40 mM), methionine (10 mM), isoleucine (30 mM), leucine (45 mM), tyrosine (10 mM), phenylalanine (16 mM), histidine (5 mM), lysine (15 mM), arginine (10 mM), glutamine (20 mM), asparagine (20 mM), tryptophan (20 mM), cysteine (20 mM). The amino acid concentrations are approximate concentrations that vary by lot, according to the information provided by the manufacturer. Oocytes were transferred to ND96 (pH 7.4) buffer for one hour efflux period. The concentration of free amino acids in the buffer was quantified by LC-MS: a Dionex Ultimate 3000 HPLC system by an autosampler (Thermo Fisher Scientific, CA, USA). The HPLC system was interfaced with the Exactive Plus Fourier transfer mass spectrometer with an electrospray ionization source. 1 µL of oocyte buffers were directly injected to LC-MS. The chromatography mobile phases were solvent A (0.3% formic acid in AcCN) and solvent B (AcCN/100 mM ammonium formate: 20/80). The column was developed at a flow rate of 600 μL min^−1^ with the following concentration gradient of solvent B: 20% B in 4 min, 20% B to 100% B in 10 min, hold at 100% B for 2 min, from 100% B to 20% B in 0.1 min, and finally, re-equilibrate at 2% B for 10 min. The electrospray ionization source was operated in positive ion mode. Data acquisition and analysis were performed through Xcalibur software (version 2.2). Quantification was carried out by measuring peak area relative to that corresponding to ^15^N amino acids (10 µM). Experiments were performed four times and representative results are shown.

### *B. cinerea* infections

*B. cinerea* strain B05.10 strain was used (Van Kan *et al*., 2017). B05.10 was grown on Malt Extract medium for two weeks prior to infection, washed with deionized water and filtered twice with Miracloth (0.78 microns) to a final concentration of 2.5 × 10^5^ spores/ml determined by hemocytometer counting. Spores were re-suspended in an inoculation medium composed of 0.1 M sucrose, 0.01 M KH_2_PO_4_ pH 4.58 (filter sterilized), 0.05% Tween 20. A volume of 5 µl of spores in the inoculation media was pipetted on each half leaf for all the genotypes to test for resistance. The growth chamber conditions were short day chamber (10 hours light/14 hours dark, approximately 120 μmol m^−2^ sec^−1^). The infected plants were put under the bench in the lab at 25 °C, then transferred into a plastic tray (56 × 36 × 6 cm) with the lid (56 × 36 × 18 cm). 1 L warm water (at 37 °C) was added to the tray before being sealed with tape to maintain humidity during infection. Lesion progression was monitored regularly and scored/recorded four days post-infection.

### Histochemical GUS analyses

To generate UmamiT20-GUS, the full native gene without the terminal codon and a promoter region exactly 3 kb upstream of the starting ATG was synthesized. The synthesis product was cloned into the GUS expression vector construct pUTkan using *Kpn*I and *BamH*1 sites (Pratelli *et al*., 2010). Col-0 plants were transformed using the finalized vector via the *Agrobacterium* floral dip method (Clough & Bent, 1998). Individual transformants were selected on ½ salt strength MS medium with 50 mg/mL hygromycin. For histochemical GUS analysis (to detect local induction of the UmamiT20 promoter and translation fusion), plants were stained using the GUS stain solution and followed the protocol (Yang *et al*., 2018). Infected rosette leaves were collected four days post-infection. Samples were incubated at 37 °C in GUS staining solution for 48 hours and then analyzed by light microscopy (Nikon TE3000). GUS activity was first detectable at four days post infection using two independent lines in three independent repeats with a nontransgenic wild-type Col-0 control stained in parallel (UmamiT20-GUS fusion line 1 and UmamiT20-GUS fusion line 2 in Figure 3 and Figure S1).

### Analysis of *B. cinerea* induced lesion size

Photographs of each infected leaf were analyzed in Adobe Photoshop to determine the size of each lesion in mutants and Col-0 lines. A mask was carefully drawn around the irregular area of every lesion of necrotic tissue to determine the number of pixels in each lesion and compared to a standard ruler in the same image to determine the area for each lesion square millimeters.

## Results and Discussion

### UmamiT transporter *mRNAs* accumulate during *B. cinerea* infection

To identify host amino acid transporters that may be involved in pathogenesis, we searched public data sets for *UmamiT* mRNAs that increased during infection of *Arabidopsis* by *B. cinerea* (Birkenbihl *et al*., 2012; Coolen *et al*., 2016; Ferrari *et al*., 2007; Ingle *et al*., 2015; Lemonnier *et al*., 2014; Mulema & Denby, 2012; Windram *et al*., 2012; Zhang *et al*., 2013). For reference, analyses of eight public microarray or RNA-seq datasets (GEO) indicated that mRNA levels of two glucose transporters, *SWEET4* and *STP13*, with important roles in susceptibility or resistance to *B. cinerea* infection (Chong *et al*., 2014; Lemonnier *et al*., 2014) increased during infection of *Arabidopsis* leaves by *B. cinerea* (Table 1). *STP13*, which had also been shown to be important for resistance against *B. cinerea*, was consistently upregulated in seven of the eight datasets with an average 9.8x fold increase across datasets. Although *SWEET4* had previously been shown to be important for susceptibility to *B. cinerea*, only one dataset indicated an increase in mRNA levels, five observed no significant change, and two did not evaluate this transcript (Lemonnier *et al*., 2014; Chong *et al*., 2014). Out of the UmamiTs paralogs analyzed, *UmamiT18* mRNA levels were increased in four of the eight studies, with an average 2.6x fold increase, while *UmamiT20* mRNA levels were up in two experiments, with an average 3x increase (Table 1). *UmamiT18*, also known as *Siliques Are Red1* (*SIAR1)*, had previously been shown to be a key player for amino acid translocation from leaves to growing siliques (Ladwig *et al*., 2012). The combination of multiple different inoculation buffers and different fungal stains used may explain the variability among these studies. We chose *UmamiT18* (AT1G44800) and *UmamiT20* (AT4G08290) as candidates for further evaluation (Table 1; Fig 1).

**Fig 1.**
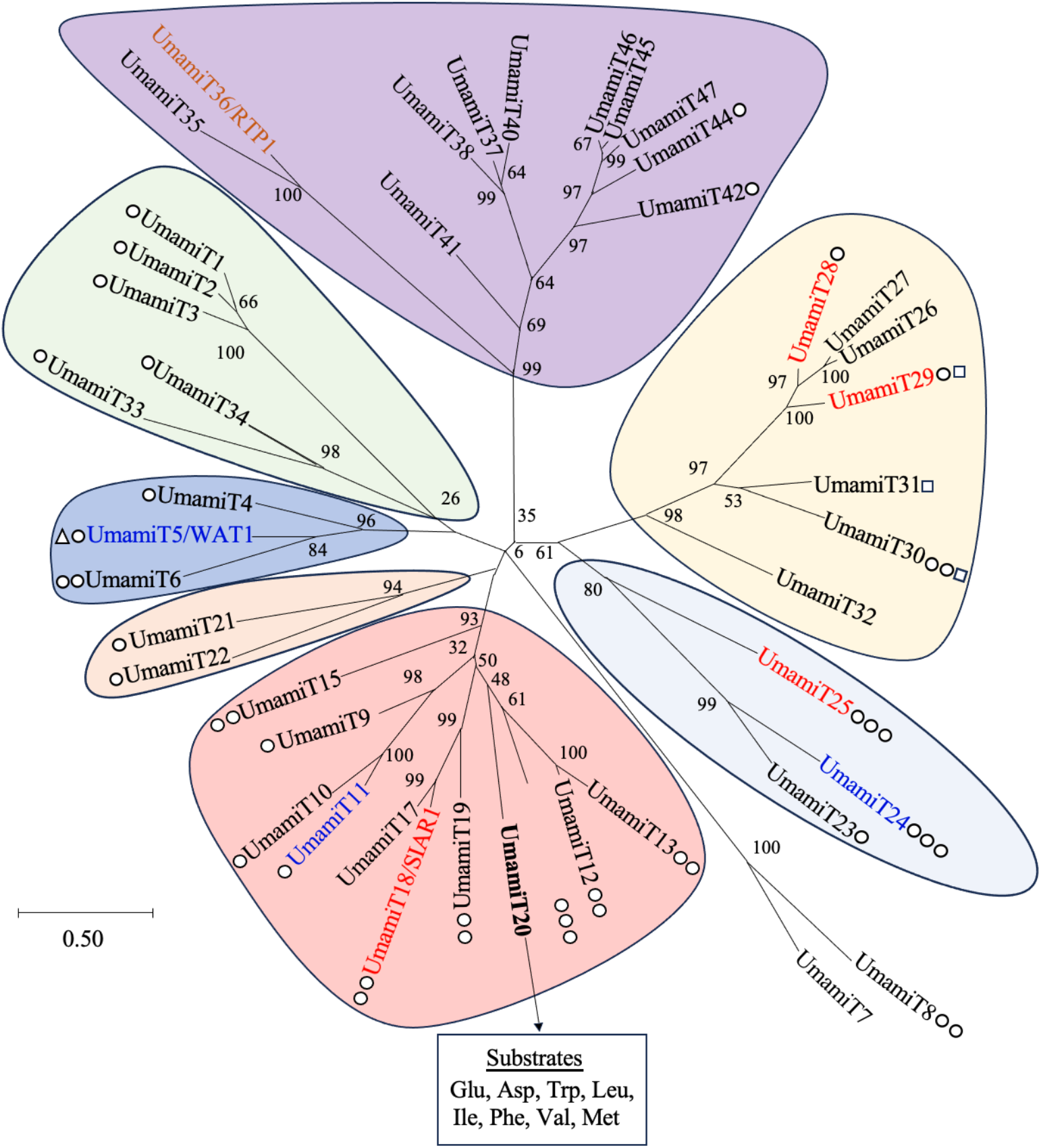
Phylogenetic family tree for *Arabidopsis* UmamiTs. Phylogenetic evolutionary history was inferred by using the Maximum Likelihood method based on the Le & Gascuel model using 1000 bootstraps (Le & Gascuel, 2008). The highly variable N- and C-terminal regions of the UmamiT proteins were not included. The tree was drawn to scale, with branch lengths corresponding to the number of substitutions per site. The analysis involved 45 amino acid sequences. Subcellular localization: plasma membrane (red), tonoplast membrane (blue), or ER membrane (dark orange). White circles indicate the number of reported amino acid substrates: 1-5 (single circle), 6-10 (two circles), >10 (three circles) while a white triangle indicates a confirmed auxin transporter and white squares indicate glucosinolate substrates. Evolutionary analyses were conducted in MEGA X (Kumar *et al*., 2018).

### UmamiT20 is a functional amino acid transporter

UmamiT20 is closely related to UmamiT18, also known as *Siliques Are Red1* (SIAR1), which had previously been shown to be a key player for amino acid translocation from leaves to growing siliques (Ladwig *et al*., 2012; Kim *et al*., 2021). Other UmamiT family members, such as UmamiT11, 14 and 29 were shown to function as amino acid transporters in transport of a broad range of amino acids when expressed in *Xenopus* oocytes or yeast (Ladwig *et al*., 2012; Müller *et al*., 2015; Besnard *et al*., 2016; Besnard *et al*., 2018; Zhao *et al*., 2021). We utilized machine and deep learning-based prediction models to explore the predicted substrate interactions of UmamiT20 and the predicted K_m_ with L-amino acids (Table 2) (Kroll *et al*., 2021; Kroll *et al*., 2023). According to the *SPOT* transporter-substrate pair prediction model, glutamine, one of the most abundant amino acids in plants, was predicted as substrate for UmamiT20 with a prediction score of 0.86. This score was higher than that of the known glutamine transporter UmamiT18 (*SPOT* prediction score: 0.83) (Kroll *et al*., 2023) (Table 2). To determine amino acid transport experimentally, UmamiT20 expressing *Xenopus* oocytes were injected with a ^15^N-labelled amino acid mix and the amount of free amino acids in the buffer, exported by UmamiT20, was quantified by LC-MS. UmamiT20 expressing oocytes were capable of exporting glutamine, isoleucine, leucine, valine, methionine, phenylalanine, tryptophan, and asparagine, indicate that UmamiT20 functions as a broad-spectrum amino acid efflux transporter (Fig. 2).

**Table 2.** Prediction of protein / metabolite pairing and K_M_ predictions using SPOT and K_M_ machine learning-based prediction tools for UmamiT20 and L-amino acids (https://deepmolecules.org/) (Kroll *et al*. 2021; Kroll *et al*. 2023). Substrates validated by transport assays in *Xenopus* oocytes in this study are marked in bold.

|  | Prediction<br>Score | K<sub>m</sub><br>prediction<br>[mM] | KEGG<br>compound<br>entry |
| --- | --- | --- | --- |
| **Glutamine** | 0.86 | 0.62 | C00064 |
| **Asparagine** | 0.53 | 0.50 | C00152 |
| **Tryptophan** | 0.21 | 0.16 | C00078 |
| **Leucine** | 0.14 | 0.23 | C00123 |
| **Isoleucine** | 0.13 | 0.97 | C00407 |
| **Phenylalanine** | 0.09 | 0.21 | C00079 |
| **Valine** | 0.070 | 1.44 | C00183 |
| **Methionine** | 0.02 | 0.93 | C00073 |
| --- | --- | --- | --- |
| Proline | 0.34 | 0.90 | C00148 |
| Alanine | 0.21 | 1.73 | C00041 |
| Tyrosine | 0.16 | 0.30 | C00082 |
| Glycine | 0.16 | 1.63 | C00037 |
| Glutamate | 0.12 | 1.27 | C00025 |
| Threonine | 0.12 | 1.69 | C00188 |
| Proline | 0.24 | 0.64 | C16435 |
| Serine | 0.35 | 0.92 | C00716 |
| Threonine | 0.12 | 1.69 | C00188 |
| Tyrosine | 0.21 | 0.22 | C01536 |

**Fig 2.**
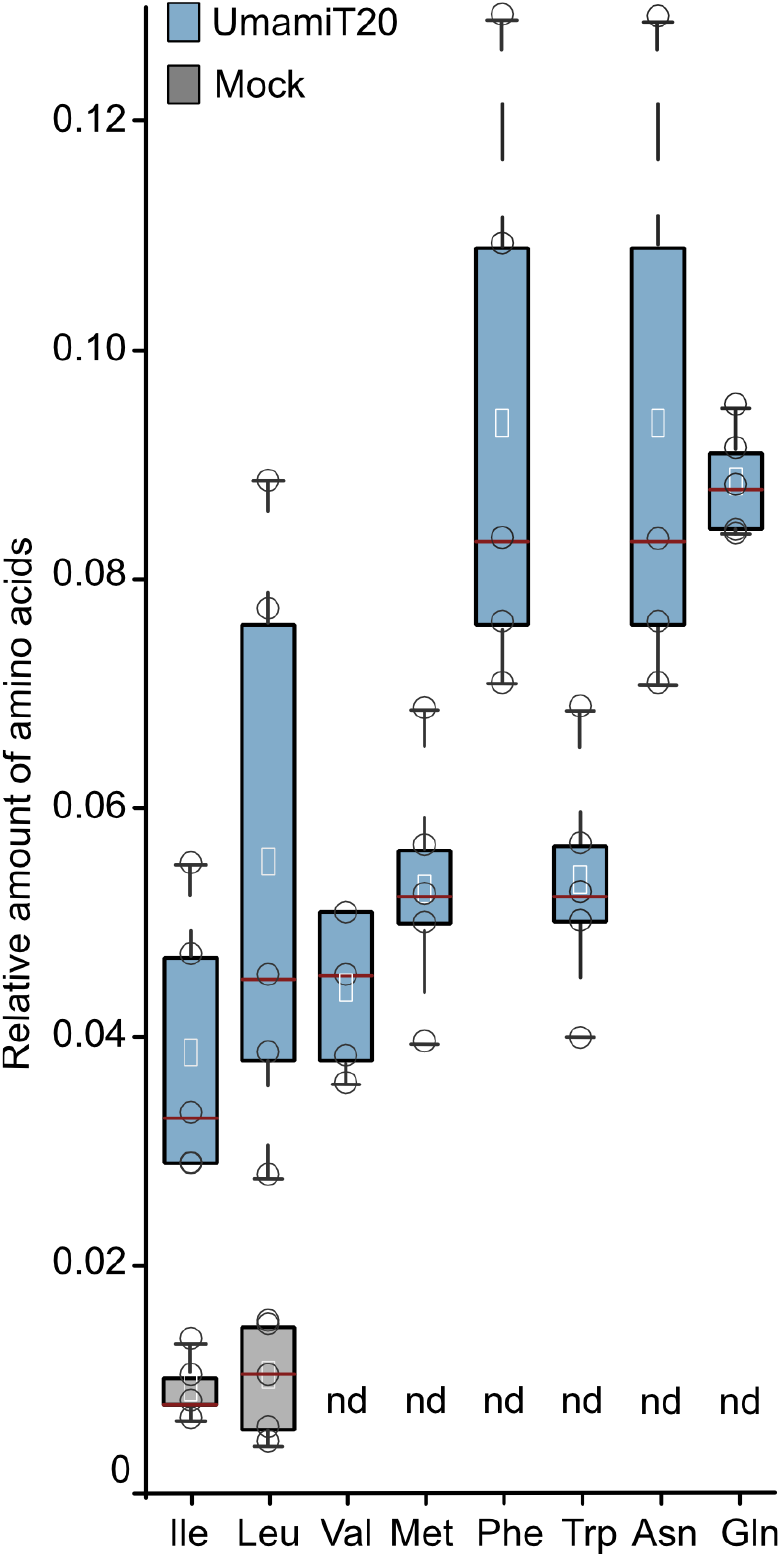
UmamiT20 functions as a broad specificity amino acid transporter. *Xenopus* oocytes injected with *UmamiT20* cRNA (light blue) and water as control (gray) were injected with 50 nL of ^15^N labeled amino acid mixture. Efflux was measured by quantifying the concentration of free amino acids in the buffer. Each circle represents an independent measurement. The lines of boxes represent the 25^th^ percentile (top) and 75^th^ percentile (bottom) respectively. Red line indicates the median. ND = not detected, n = 5 ± SE.

### UmamiT20 accumulates in the leaf vasculature close to the site of infection

To determine whether UmamiT20 accumulates at infection sites, and to obtain insights into the spatial and temporal protein accumulation during *Botrytis* infection, we characterized *Arabidopsis* lines expressing translational UmamiT20-GUS fusions driven by their own promoter. We did not observe UmamiT20-GUS protein accumulation during the initial stages of the infection (first three days), however clear accumulation four days after infection in the leaf vasculature, a time at which substantial necrosis was evident (Fig. 3; Fig. S1). GUS reporter activity was detectable specifically in the vasculature surrounding the sites of the infection in three independent biological replicates using two independent transformants. By contrast, GUS reporter activity was not observed in any of the mock controls (Fig. S1). We concluded that the abundance of UmamiT20-GUS protein increases in the vasculature during later stages of infection.

**Fig 3.**
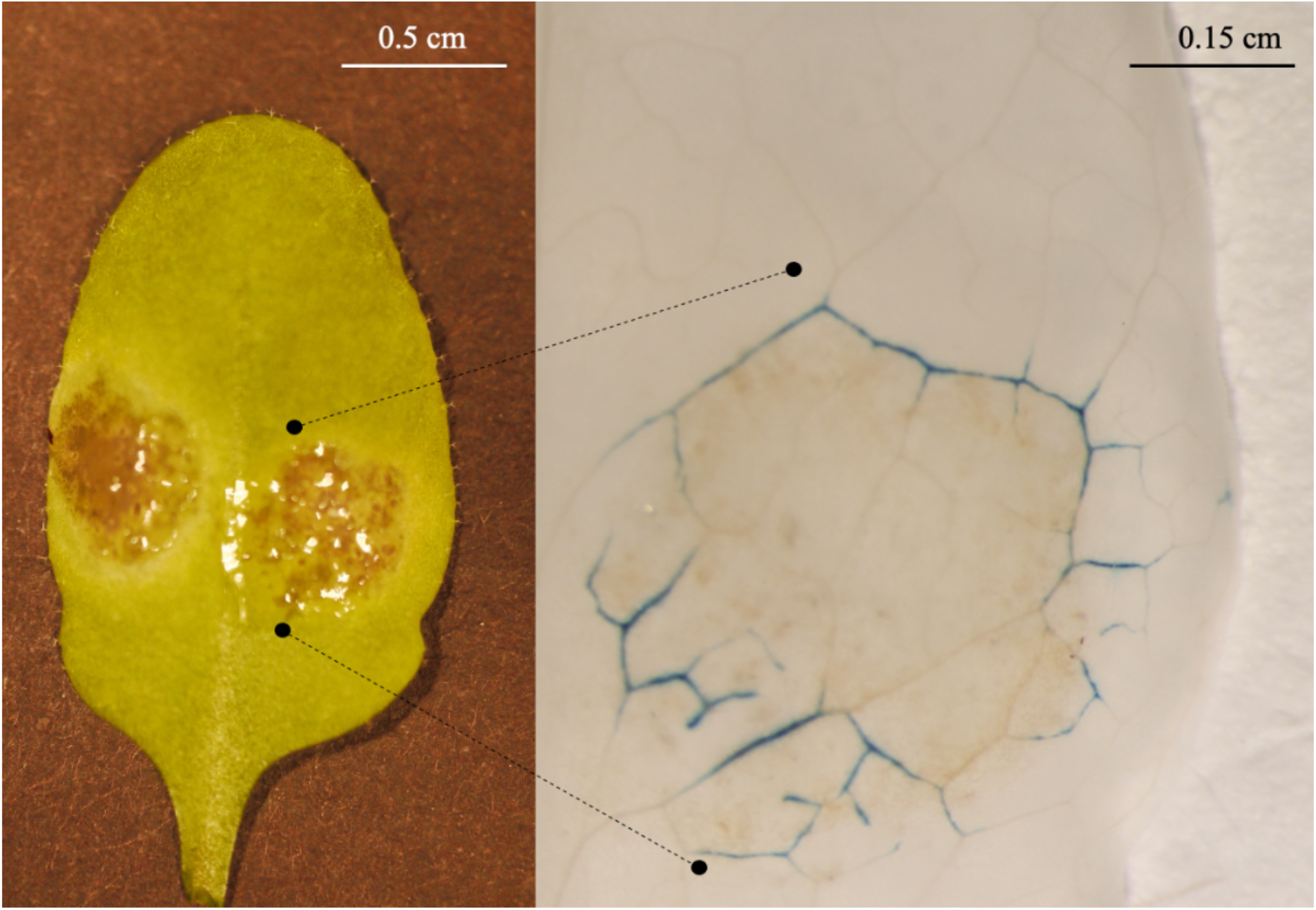
UmamiT20-GUS accumulation in veins surrounding *B. cinerea* caused lesions. Images were taken four days post infection (left) and magnified post GUS staining (blue product; right) of a representative *B. cinerea* infected leaf. Shown is the result from UmamiT20-GUS fusion line 2. Comparable data were observed in 3 independent experiments for a total of 9 leaves, 1 leaf per individual plant. UmamiT20-GUS line 1 displayed comparable induction patterns. Additional images available in Fig. S2.

### UmamiT20 localizes to the plasma membrane in *N. benthamiana*

Previously, UmamiT14 and UmamiT18 were to localized to the plasma membrane in *Arabidopsis* (Ladwig *et al*., 2012), similarly UmamiT11, UmamiT14, UmamiT28, and UmamiT29 had been shown to localize to the plasma membrane in *N. benthamiana* (Müller *et al*., 2015). Many of the UmamiTs characterized so far function as plasma membrane transporters, while WAT1 can function as a vacuolar auxin transporter (Ranocha *et al*., 2013). To determine the subcellular localization of UmamiT20, a translational UmamiT20-eGFP fusion was expressed transiently in *N. benthamiana* leaves. In three independent biological repeats, UmamiT20-eGFP derived fluorescence was detectable on the peripheral side of chloroplasts, indicative of predominant plasma membrane localization (Fig. 4) and consistent with a role of UmamiT20 in amino acid uptake or release in the vicinity of the infection sites.

**Fig 4.**
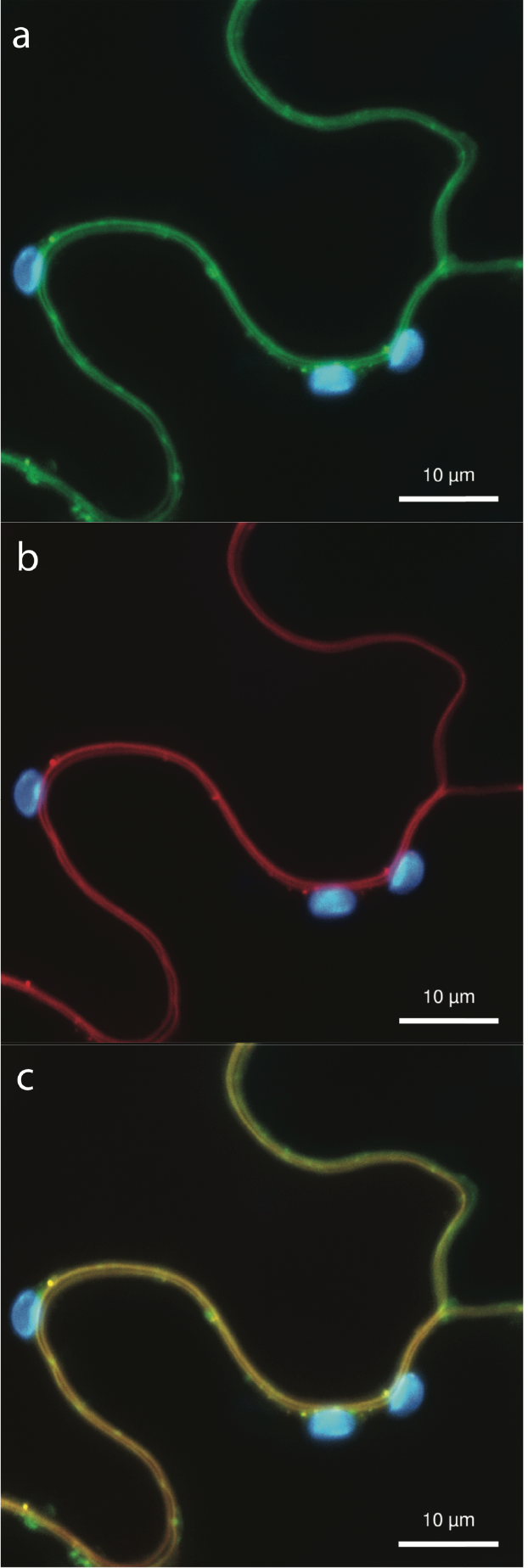
Subcellular localization of UmamiT20-eGFP fusions in tobacco leaves. Confocal images (maximal projection of a z-stack) of *Agrobacterium*-infiltrated *N. benthamiana* leaf cells. ZmSWEET13a:mCherry was used as a plasma membrane marker. The eGFP signal from panel a (522–572 nm) (green) and mCherry fluorescence from panel b (600-625 nm) (red) were merged with fluorescence derived from chloroplasts (667–773 nm) (blue). (a) UmamiT20:eGFP (C-term) with chloroplast fluorescence; (b) ZmSWEET13a:mCherry (C-term) with chloroplast fluorescence; (c) Merge of eGFP, mCherry, and chloroplast fluorescence. Fluorescence was visualized using confocal laser scanning microscopy 3 days after *Agrobacterium* infiltration, 12-16 hours after induction of UmamiT20-GFP with ß-estradiol. Cells with high expression levels showed fluorescence in puncta along the plasma membrane.

### *umamiT20* mutants show reduced disease symptoms

Since UmamiT20 mRNA and protein accumulated late in *B. cinerea* infection (Table 1), we explored whether pathogenicity is affected in *Arabidopsis* knock-out mutants. The T-DNA insertion mutant *umamit20-1* is an apparent null mutant as judged by the absence of detectable cDNA using PCR amplification (Fig. S3). *umamit20-1* homozygous mutants showed decreased susceptibility to *B. cinerea* in five independent biological replicates compared to the wild-type Col-0 (Fig. 5; Fig S2). Phenotypic quantitation by image analysis showed that lesions on *umamit20-1* leaves were ∼36% on average relative to Col-0 (Fig. 5). CRISPR/Cas9 was used to generate an independent knock-out line, *umamit20-2* (Fig S4). Five independent infection assays with both the T-DNA insertion and CRISPR-Cas9 mutants showed a decrease in lesion size relative to wild-type controls (Fig. 5). The lesion size for the *umamit20-2* CRISPR-Cas9 knock-out line was ∼30% in size on average compared Col-0. In contrast, the *umamit18-1* and *umamit18-2* mutant lines (Ladwig *et al*., 2012), showed no significant phenotypic difference regarding disease symptoms by *B. cinerea*. (Fig S5). Thus UmamiT20 functions as a host susceptibility gene, analogous to SWEET4 (Chong *et al*., 2014), while UmamiT18 does not. Of note, two other studies had also invoked members of this family in pathogen susceptibility: WAT1 (UmamiT5; At1g75500) and RTP1 (UmamiT36; At1g70260) (Denancé *et al*., 2013; Ranocha *et al*., 2013; Pan *et al*., 2016). Both belong to more distantly related clades of the UmamiT family (Fig. 1). WAT1 was shown to function as a vacuolar auxin transporter, while RTP1 had been suggested to negatively affect host resistance, in particular to biotrophic pathogens, but not to the necrotrophic *B. cinerea*, possibly by affecting signaling processes that control ROS production, cell death and *PR1* gene expression. UmamiT29, Umamit30, and UmamiT31 have recently been reported to transport glucosinolates which play a protective role against pathogens (Xu *et al*., 2023; Meyer *et al*., 2023).

**Fig 5.**
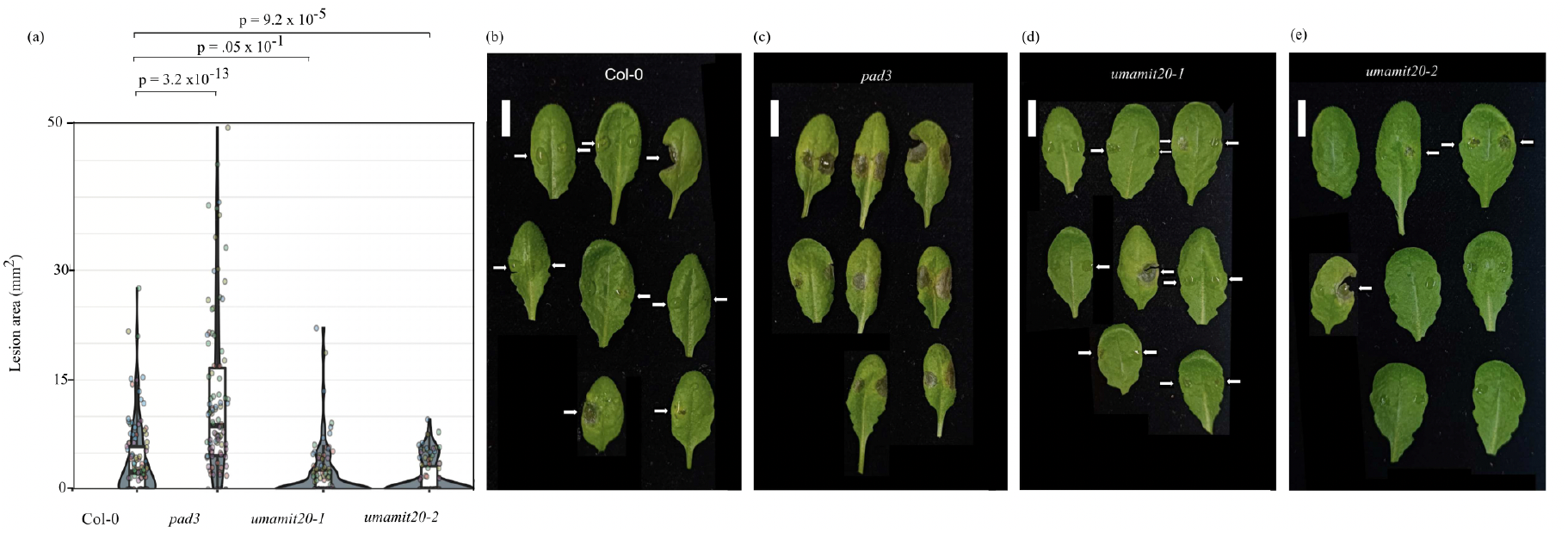
*B. cinerea* infection of Col-0 and *umamit20-1* and *umamit20-2* mutants. (a) The plots of lesion size in Col-0 compared to *umamit20-1*, and *umamit20-2* and *pad3*, a highly susceptible control. The data shown are for experiments were executed for five independent replicates. Note that in one of the five replicates used 10 μL volume of inoculum on each half leaf (standard assay volume was 5 μL). Each circle is a single lesion from a droplet inoculation. The p-values are shown using Kruskall/Wallis-Wilcoxon test, FDR: 0.05 as the data are not normally distributed, a non-parametric statistical analysis was used. (b, c, d, e) Representative images of infected leaves from a single replicate from each genotype. Scale bars: 1 cm.

Similar as for the role of SWEETs in susceptibility, we invoke two non-exclusive hypotheses when UMAMITs are activated: overcoming “pathogen nutrient starvation” or “nutrient triggered immunity” (Bezrutczyk *et al*., 2018b; Prior *et al*., 2021; Tünnermann *et al*., 2022). The “pathogen starvation” hypothesis proposes that the decreased host susceptibility in the *umamit20* mutants is due to insufficient supply with organic nitrogen resulting in the need of activating a transporter. Interestingly, recent evidence suggests that *B. cinerea* may act as a sink for host amino acids during infection. When infected by *B. cinerea*, sunflower cotyledons showed a significant decrease in amino acid levels while radiolabeled amino acids increased in the fungus (Dulermo *et al*., 2009). The translocation of amino acids from host to pathogen requires efflux transport mechanisms in the host and importers in the fungus. UmamiT20 could serve as a host efflux transporter for certain amino acids that contribute to the nutrition of fungal growth and reproduction. However, since UmamiT20 accumulates in the vasculature and not in the cells surrounding the hyphae, UmamiT20 may be involved in delivering or removing amino acids from other plant tissues and organs.

The second hypothesis, “nutrient triggered immunity”, proposes that secreted nutrients act as signals that trigger host defense responses (Gebauer *et al*., 2017). Several studies support the nutrient signaling hypothesis. *Arabidopsis sweet11/12* double mutants accumulated sugars that could prime salicylic acid-based immune responses, responsible for reduced susceptibility to the fungus *Colletotrichum higginsianum* (Gebauer *et al*., 2017; Biemelt & Sonnewald, 2006). Similarly, ectopic overexpression of the *Cationic Amino Acid Transporter 1* (CAT1), or suppression of the *Arabidopsis Lysine Histidine Transporter 1* (LHT1) led to activation of salicylic acid-based immune responses and increased resistance (Yang & Ludewig, 2014). In rice, pre-treatment of leaves with glutamate resulted in a concentration-dependent increase in resistance to the fungus *Magnaporthe oryzae* and triggered induction of immunity related genes in leaves and roots (Kadotani *et al*., 2016). UmamiT20 could thus contribute to both pathogen nutrition and host nutrient-activated immune signaling, Furthermore, the comparison of the amino acid substrates of UmamiT20 and the metabolic requirements of the pathogen show clear overlap. *B. cinerea* can use seven of the eight amino acids substrates of UmamiT20 as an N source on synthetic media, all substrates except valine (Wang *et al*., 2018).

Future studies using *sweet4*/*umamit20* double mutants may help determine whether the loss of multiple transporters quantitatively increases the resistance phenotype and helps to evaluate whether nutrient availability affects disease outcome (Biemelt & Sonnewald, 2006). Further research could investigate whether the combinatorial loss of SWEET4 and UMAMIT20 activity alters hormone levels, such as salicylic acid or jasmonic acid, involved in immune signaling. Conversely, the loss of multiple nutrient transporters could negatively affect the energy mobilization required for the host’s immune response, thereby increasing susceptibility. To understand the fungal amino acid transporters that might play a role in the interaction between the host and the pathogen, we examined the *B. cinerea* B05.10 genome for putative fungal amino acid transporters (compared to known yeast amino acid facilitators) and identified 27 candidate genes for further evaluation (Bianchi *et al*., 2019; Table S2). Future work to untangle these competing explanations and understand the precise role of each of the host transporters during pathogenesis is an area of great interest.

## Conclusions

In this report, we demonstrate that UmamiT20 functions as amino acid transporter and demonstrate it’s necessary during *B. cinerea* infection in *Arabidopsis*. The reduced susceptibility in *umamiT20* mutant lines intimates that amino acids levels in the infected leaf may be important to determining the outcome of host pathogen interactions. This study introduces the *UmamiT20* amino acid transporter gene as a new susceptibility gene, possibly implicating *UmamiT* gene family members in necrotrophic host-pathogen systems. Combinatorial CRISPR-based mutagenesis of *SWEET* and *UmamiT* genes in crop plants may offer new ways to combat *B. cinerea* in crops such as grape in the future.

## Acknowledgements

We thank Alexander Davis and Davide Sosso for constructive advice and discussions. This work was supported by grants from the National Science Foundation (IOS-1936492); Deutsche Forschungsgemeinschaft (DFG, German Research Foundation) - Collaborative Research Center SFB1535, project ID 458090666/CRC1535/1; Deutsche Forschungsgemeinschaft (DFG, German Research Foundation) under Germany’s Excellence Strategy – EXC-2048/1 – project ID 390686111, Deutsche Forschungsgemeinschaft (DFG, German Research Foundation), DFG, project ID 391465903/GRK 2466), the Alexander von Humboldt Professorship (WBF). Work in HLJ’s lab was supported by the National Institutes of Health (R01 GM093008), the National Science Foundation (IOS-1557812) and an AES-CE Award (PPA-7517H) awarded to HJ. Work of JYK’s lab was supported by Basic Science Research Program through the National Research Foundation of Korea (NRF) funded by the Ministry of Education (No. NRF-2019R1A6A1A10073079). Work in GP’s lab was supported by the National Science Foundation (MCB-1519094), the Virginia Agricultural Experiment Station and the Hatch Program of the National Institute of Food and Agriculture, U.S. Department of Agriculture (VA-135908).

## Author contributions

MJP, JYL, FL, HBY, QC, CD prepared or/and performed infection assays. MJP and HBY performed GUS assays and bioinformatic analysis. JYK, DW, KK performed efflux assays in oocytes. MB performed GFP fusion localization assay. DW and CD generated *umamit20-2* CRISPR-Cas9 mutant, MJP, JYK, JYL, QC, CZ, LQC, MCJ, GP, HJ, WBF performed experimental design and wrote the manuscript.

## Supplementary Information

**Table S1:** Compilation of primer sequences used in this study.

**Table S2:** Candidate amino acid transporters in *B. cinerea* B05.10 strain

**Figure S1**. Additional images of two translational UmamiT-GUS fusion lines during infection.

**Figure S2**. Additional images of *umamit20* knock-out or wild-type plants during infection.

**Figure S3**. Schematic representation of the T-DNA insertion line (*umamit20-1*) used in this study.

**Figure S4**. Schematic representation of the CRISPR/Cas9 line (*umamit20-2*) used in this study.

