## Supplement for "The *Arabidopsis* amino acid transporter UmamiT20 confers *Botrytis cinerea* susceptibility"

The following Supporting Information is available for this article:

**Table S1:** Compilation of primer sequences used in this study.
**Table S2:** Candidate amino acid transporters in *B. cinerea* B05.10 strain
**Figure S1.** Additional images of two translational UmamiT-GUS fusion lines during infection.
**Figure S2.** Additional images of *umamit20* knock-out or wildtype plants during infection.
**Figure S3.** Schematic representation of the T-DNA insertion line (*umamit20-1*) used in this study.
**Figure S4.** Schematic representation of the CRISPR/Cas9 line (*umamit20-2*) used in this study.
**Figure S5.** Images and data of *umamit18* mutants or wildtype plants during infection.

**Table S1.** Compilation of primer sequences used in this study.

| Primer name | Primer Sequence: | Purpose |
|---|---|---|
| ***UmamiT20* ORF fw** | 5’ GACAAGTTTGTA CAAAAAAGCAGGCTCATCAATAACCATGGAG GGTGTTAGT 3’ | Amplification of *UmamiT20* ORF for *N. benthamiana* expression |
| ***UmamiT20* ORF rev** | 5’GACCACTTTGTACAAGAAAGCTGGGTACTAT ATTCCAGATGTGTTACTATGCGC 3’ | Amplification of *UmamiT20* ORF for *N. benthamiana* expression |
| **DW185** | 5’ggctacggtctctattgGTGCAACTATGCATAAGTTAG TTTAGAGCTAGAAATAG 3’ | *UmamiT20*-target1+BasI fw |
| **DW186** | 5’ggctacggtctctaaacGTTCTGGCCTTGGTTAAGAGT GACCAATGTTGCTCCCTC 3’ | *UmamiT20*-target 2 + BsaI rev |
| **MR9** | 5’GATTGAATCCTGTTGCCG 3’ | Colony PCR |
| **MQ5 fw** | 5’CGTAGCTAACAATAGACGTG 3’ | *UmamiT20* genotyping primer |
| **MQ5 rev** | 5’GCTGATGTCATGTTCATCC 3’ | *UmamiT20* genotyping primer |
| **MQ5 seq** | 5’ACTCTACATGACTCTCGAT 3’ | *UmamiT20* sequencing primer |

**Table S2.** Candidate amino acid transporters in *B. cinerea* B05.10 strain. A thorough search of the annotated B05.10 genome compared to known yeast amino acid transporters resulted in the 27 putative candidates for *B. cinerea* amino acid transporters in the Joint Genome Institute (JGI) database (https://img.jgi.doe.gov) (Bianchi *et al.*, 2019; Van Kan *et al.*, 2017).

| ***B. cinerea* JGI Gene ID** | **Current Gene Annotation** | **Putative Transporter Family** |
|---|---|---|
| **2973936300** | Amino acid permease | YAT family |
| **2973917861** | Amino acid permease | YAT family |
| **2973939629** | Amino acid permease | YAT family |
| **645752960** | Amino acid transporter | YAT family |
| **645764320** | Amino acid transporter | YAT family |
| **645760795** | Amino acid transporter | YAT family |
| **645752127** | Amino acid transporter | YAT family |
| **645764101** | Amino acid transporter | YAT family |
| **645764321** | Amino acid transporter | YAT family |
| **645753521** | Amino acid transporter | YAT family |
| **645758843** | Amino acid/polyamine/organocation transporter | APC family |
| **645754001** | Amino acid/polyamine/organocation transporter | APC family |
| **645757575** | Amino acid/polyamine/organocation transporter | APC family |
| **645759911** | Amino acid/polyamine/organocation transporter | APC family |
| **645764102** | Aromatic amino acid and leucine permease | YAT family |
| **645761693** | Aspartate/glutamate transporter | YAT family |
| **645767906** | General amino acid transporter | YAT family |
| **645766592** | General amino acid transporter | YAT family |
| **645763806** | General amino acid transporter | YAT family |
| **645762569** | Lysine/proton symporter | AAT family |
| **645765536** | Methionine transporter | LAT family |
| **645765505** | Methionine transporter | LAT family |
| **645754377** | Proline transporter | YAT family |
| **645756626** | Proline transporter | YAT family |
| **645755103** | Aspartate/glutamate transporter | Solute carrier family 25 |
| **645756292** | Vesicular inhibitory amino acid transporter | Solute carrier family 32 |
| **645766422** | Proton-Coupled amino acid transporter | Solute carrier family 36 |

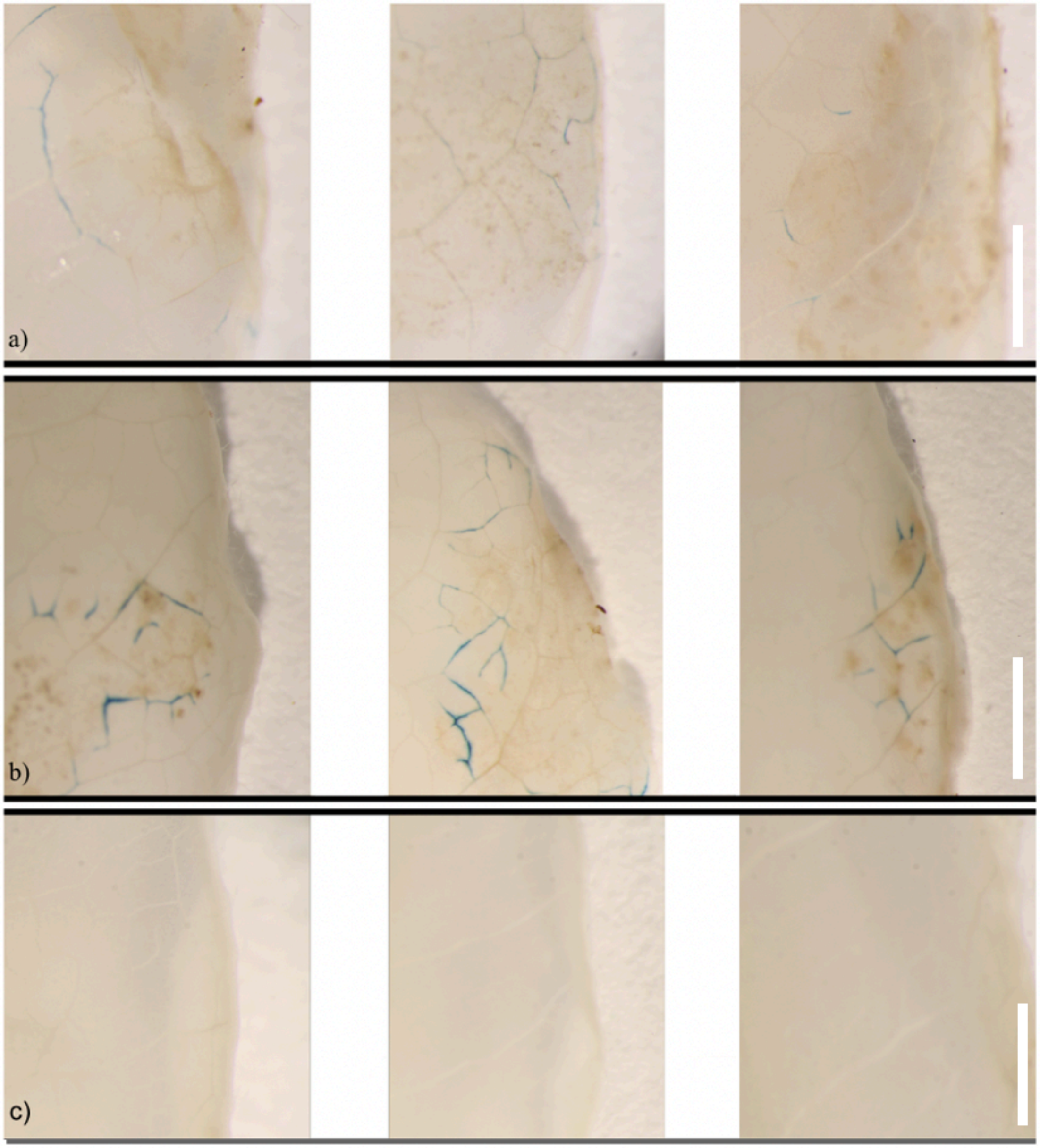


**Fig S1.** Representative images of UmamiT20-GUS fusion lines. (a) Images from heterozygous UmamiT20-GUS fusion line 1, 80% of plants displayed UmamiT20-GUS accumulation (n=15 plants). (b) Images from homozygous UmamiT20-GUS fusion line 2, 100% of plants showed UmamiT20-GUS accumulation also in vasculature (n=9 plants). (c) Mock control with inoculation buffer alone placed on leaves from UmamiT20-GUS fusion lines 1 and 2. Scale bar: 0.25 cm.

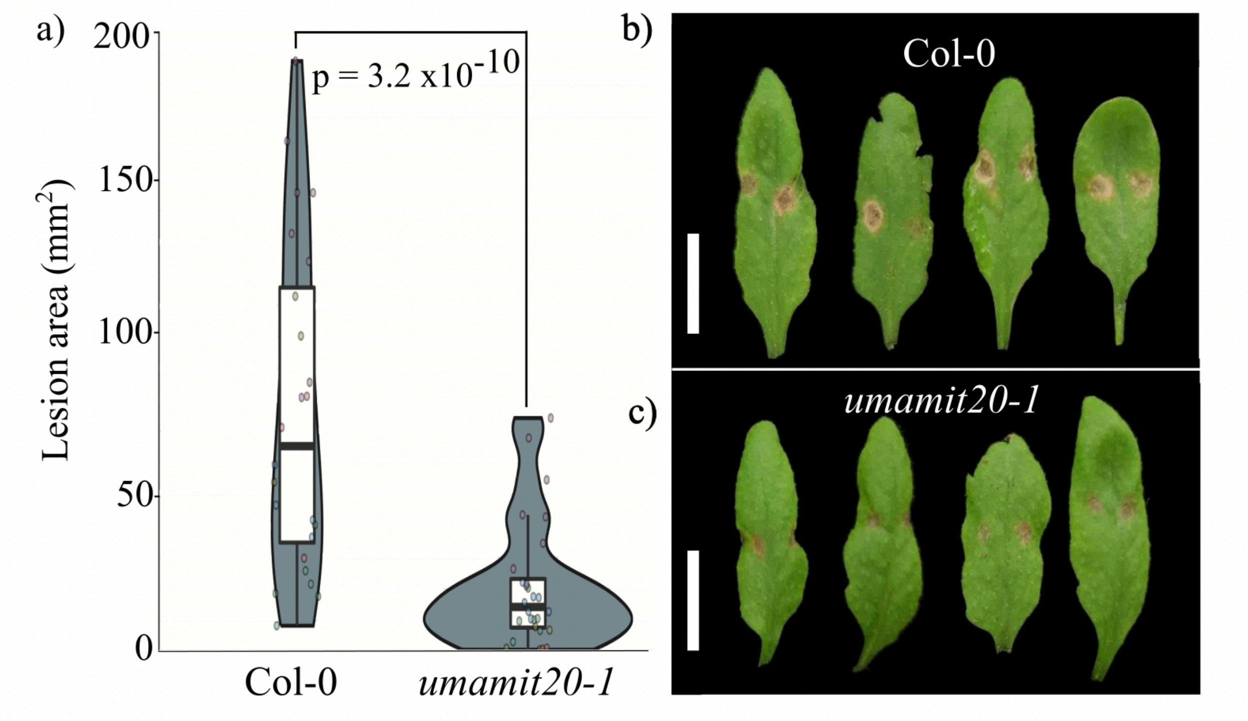


**Fig S2.** Additional data and representative images of symptoms caused by *B. cinerea* infection on leaves of homozygous *umamit20-1 knock-out* lines and Col-0 controls. (a) Plot displaying lesion sizes of wild-type Col-0 and *umamit20-1* T-DNA insertion mutant caused by *B. cinerea* infection. Infection assays were conducted over seven independent replicates. (b) Representative image of infected leaves. All leaves were collected four days after inoculation. Scale bar: 1 cm.

*UmamiT20*

*umamit20-1*   *Salk_076704*

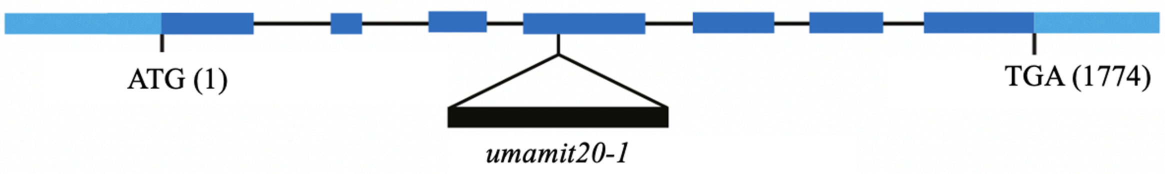


**Fig. S3** Schematic representation of the *umamit20-1* T-DNA insertion line (Salk_076704) used in this study. The light blue bars represent the 5'UTR and 3'UTR from left to right. Dark blue bars represent the exons. The insert of interest was confirmed to be in the fourth exon by PCR genotyping.

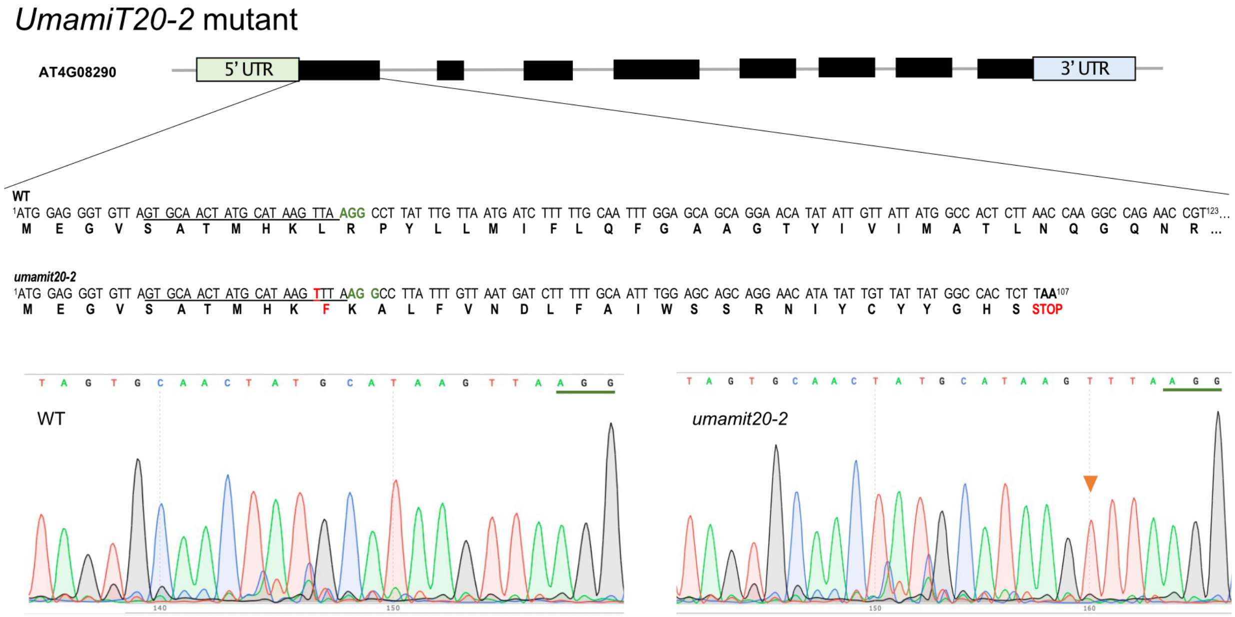


**Fig. S4** Schematic representation of the mutation in the *umamit20-2* CRISPR-Cas9 mutant

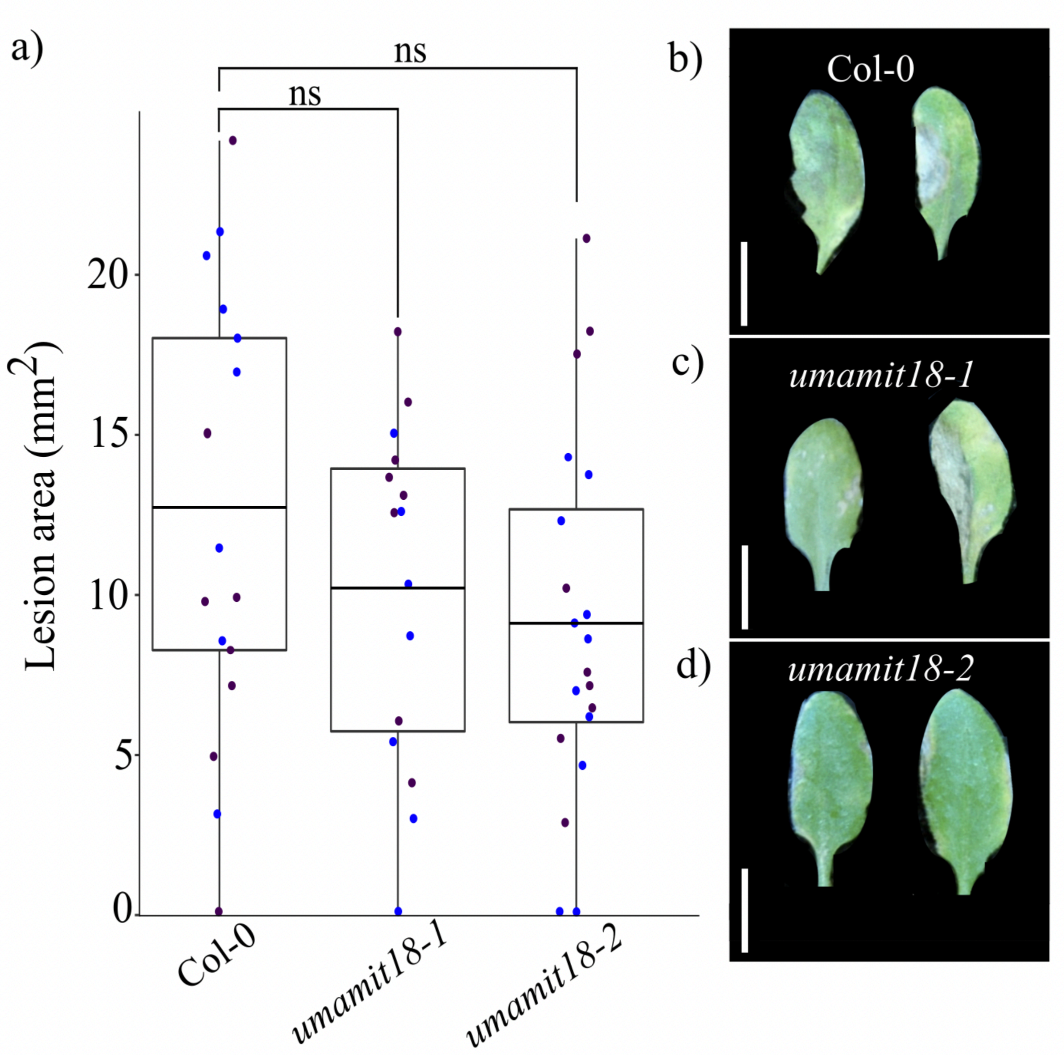


**Fig. S5**. Symptoms caused by *B. cinerea* infection on leaves of homozygous *umamit18-1 and umamit18-2* mutant lines and wild type Col-0 control lines. (a) Box and dot plot of two independent replicates of the three genotypes during infection. An ANOVA statistical test was used as the dataset was normally distributed with homogeneous variance. The p-value for Col-0 compared to *umamit18-1* is 0.48, for Col-0 compared to *umamit18-2* is 0.18. (b,c,d) Representative images of infected leaves. All leaves were collected four days after inoculation. Scale bar: 1 cm.
